# Widespread germline genetic heterogeneity of human ribosomal RNA genes

**DOI:** 10.1101/2021.07.21.453267

**Authors:** Wenjun Fan, Eetu Eklund, Rachel M. Sherman, Hester Liu, Stephanie Pitts, Brittany Ford, NV Rajeshkumar, Marikki Laiho

## Abstract

Polymorphism drives survival under stress and provides adaptability. Genetic polymorphism of ribosomal RNA (rRNA) genes derives from internal repeat variation of this multicopy gene, and from interindividual variation. A considerable amount of rRNA sequence heterogeneity has been proposed but has been challenging to estimate given the scarcity of accurate reference sequences. We identified four rDNA copies on chromosome 21 (GRCh38) with 99% similarity to recently introduced reference sequence KY962518.1. Pairwise alignment of the rRNA coding sequences of these copies showed differences in sequence and length. We customized a GATK bioinformatics pipeline using the four rDNA loci, spanning a total 145 kb, for variant calling. We employed whole genome sequencing (WGS) data from the 1000 Genomes Project phase 3 and analyzed variants in 2,504 individuals from 26 populations. Using the pipeline, we identified a total of 3,790 variant positions. The variants positioned non-randomly on the rRNA gene. Invariant regions included the promoter, early 5’ ETS, 5.8S, ITS1 and certain regions of the 28S rRNA, and large areas of the intragenic spacer. 18S rRNA coding region had very few variants, while a total of 470 variant positions were observed on 28S rRNA. The majority of the 28S rRNA variants located on highly flexible human-expanded rRNA helical folds ES7L and ES27L, suggesting that these represent positions of diversity and are potentially under continuous evolution. These findings provide a genetic view for rRNA heterogeneity and raise the need to functional assess how the 28S rRNA variants affect ribosome functions.

## INTRODUCTION

Ribosomes translate mRNA templates to proteins. The human ribosome is a large protein-RNA complex that is composed of a large 60S subunit consisting of 28S, 5S and 5.8S ribosomal RNAs (rRNA) and 47 proteins and a small 40S subunit with 18S rRNA and 33 proteins (Anger et al. 2013; Khatter et al. 2015). The rRNAs are transcribed by RNA polymerases I (5.8S, 18S, 28S rRNAs) and III (5S rRNA). RNA polymerase I (Pol I) transcribes a long polycistronic 47S rRNA precursor that is processed and cleaved into the mature 5.8S, 18S and 28S rRNAs assisted by a multitude of rRNA biogenesis proteins and small nucleolar non-coding RNAs (Moss et al. 2007). The processing, folding and maturation requires also extensive posttranslational modification of the rRNAs (Sloan et al. 2017).

Ribosomes are highly abundant in cells and their numbers reflect the translation rates of the host. However, this dependency may be nonlinear leading to defects and diseases states related to alterations in ribosomal protein stoichiometry or modifications of the ribosomal proteins and rRNAs (Emmott et al. 2019; Li and Wang 2020). The ensuing ribosome heterogeneity potentially leads to functional consequences such as abnormalities observed in ribosomopathies (Mills and Green 2017) . Functionally heterogeneous ribosomes are also referred to as specialized ribosomes. Specialized ribosomes translate subsets of mRNAs and are predicted to provide a dynamic response to stress or stimuli, or occur in a cell- or tissue-specific manner (Emmott et al. 2019).

The human multicopy rRNA genes are organized in repeat arrays on five acrocentric chromosomes 13, 14, 15, 21 and 22 (Henderson et al. 1972). Copy numbers are estimated to vary between 100 – 600 per diploid genome, are unevenly distributed on the acrocentric chromosomes, and vary between individuals (McStay 2016; van Sluis et al. 2020). The rRNA gene loci contain not only the gene arrays, but are also highly repetitive due to presence of satellite repeats and other repetitive elements rendering them challenging to study and assess their genetic variability (McStay 2016). The Telomere-to-Telomere (T2T) Consortium very recently provided in-depth mapping of these previously unannotated gene arrays covering a total of 10 Mb rDNA sequence derived from a functionally haploid CHM13 hydatidiform mole cell line . This analysis, using a combination of PacBio HiFi, Oxford Nanopore and PCR-free sequencing and bioinformatic methods, indicated chromosome-dependent variation of the rDNA copies.

While substantial progress has been made in identifying the proximal and distal sequences of the arrays, consensus or reference sequences for the human rRNA gene have been lacking (McStay 2016). Only few reference sequences have existed until recently (U13369, AL353644.34) (Gonzalez and Sylvester 1995). The reference sequence U13369.1 was deposited in 1994 (Gonzalez and Sylvester 1995) and has been used in vast majority of studies since then. Yet, reports recognizing rRNA sequence variation have been published since 1980s (Gonzalez et al. 1985; Gonzalez et al. 1988; Worton et al. 1988; Kuo et al. 1996). Recent transformation-associated recombination (TAR) cloning and long-read coding efforts, using a single chromosome 21 mouse/human hybrid as source, enabled the accurate assembly of a new 45 kb rDNA sequence, identified numerous discrepancies to the earlier reference and substantially refined it, and introduced a reference sequence (KY962518.1) which was 1.8 kb longer than the previous (Kim et al. 2018). These sequencing efforts also identified single nucleotide variants (SNV) and insertion/deletions (INDEL) in the rRNA coding and non-coding intragenic spacer (IGS) sequences based on multiple sequenced clones of chromosome 21 (Kim et al. 2018). Several earlier studies have conducted variant calling approaches of the rDNA genes, but may be compromised by calling variants representing sequencing or alignment errors due to the use of the earlier reference U13369 (Babaian 2017; Xu et al. 2017; Parks et al. 2018). The identification of variants still requires refinement and their implications are yet to be determined.

Here we identified three full and one partial rRNA gene copies on chromosome 21 (GRCh38) with 99% similarity to the most recent rDNA reference KY962518.1. We employed whole genome sequencing (WGS) data from the 1000 Genomes Project phase 3 including 2,504 individuals from 26 populations and performed variant calling against the identified rDNA sequences in GRCh38 to generate a comprehensive set of germline SNVs and INDEL variant positions (SNV/INDEL) of the rDNA arrays. We discovered a total of 3,790 variant positions on the rRNA genes. Moreover, the variants distinctly clustered on the rRNA gene coding and IGS regions, while regions for core ribosomal protein interactions and rRNA modification sites and large areas in the IGS were near invariant. A large number of variant positions, 380 SNVs and 90 INDELs, were detected on the mature 28S rRNA. The majority of the 28S rRNA variants located on the highly flexible human-expanded rRNA helical folds ES7L and ES27L. Collectively, our findings provide a genetic view for rRNA heterogeneity and suggest potential links between variants and ribosome functions.

## RESULTS

### Genomic variation of rDNA copies in GRCh38 chromosome 21

To identify rDNA loci in GRCh38 human genome reference, we performed pairwise alignment of human rDNA reference sequence assembled by using TAR cloning and long-read sequencing (GenBank: KY962518.1) against current human reference genome GRCh38. We identified three full and one partial rDNA copies on chromosome 21 (GRCh38) with 99% similarity to the reference KY962518 (Figure 1A and Table S1). The 13.3 kb rRNA coding sequence of these copies varied in length by up to 42 nt due to length variation of all but the 5.8S rRNA domain (Table 1 and Table S1). Next, we performed multiple sequence alignment of rDNA reference sequence KY962518 and the four chromosome 21 rDNA copies (Figure 1B). We identified a total of 163 SNV/INDELs among these rDNA coding regions. Over half of the variants were detected in 5’ ETS region, 15% in the 28S rRNA, 11% in 18S rRNA, whereas there were no variants in the 5.8S rRNA coding region (Table 2). Among the variants in the 28S rRNA, two 4 - 6 nt INDELs were found, which were present in two rDNA copies and absent in the other two (Figure 1C and D). The two short INDELs consisted of GC-repeats (Figure 1D), and were located in the ES7L and ES15L expansion segments (not shown). The observed variation between the repeats in one chromosome suggests that diversity between different copies is likely to exist.

**Table 1.**
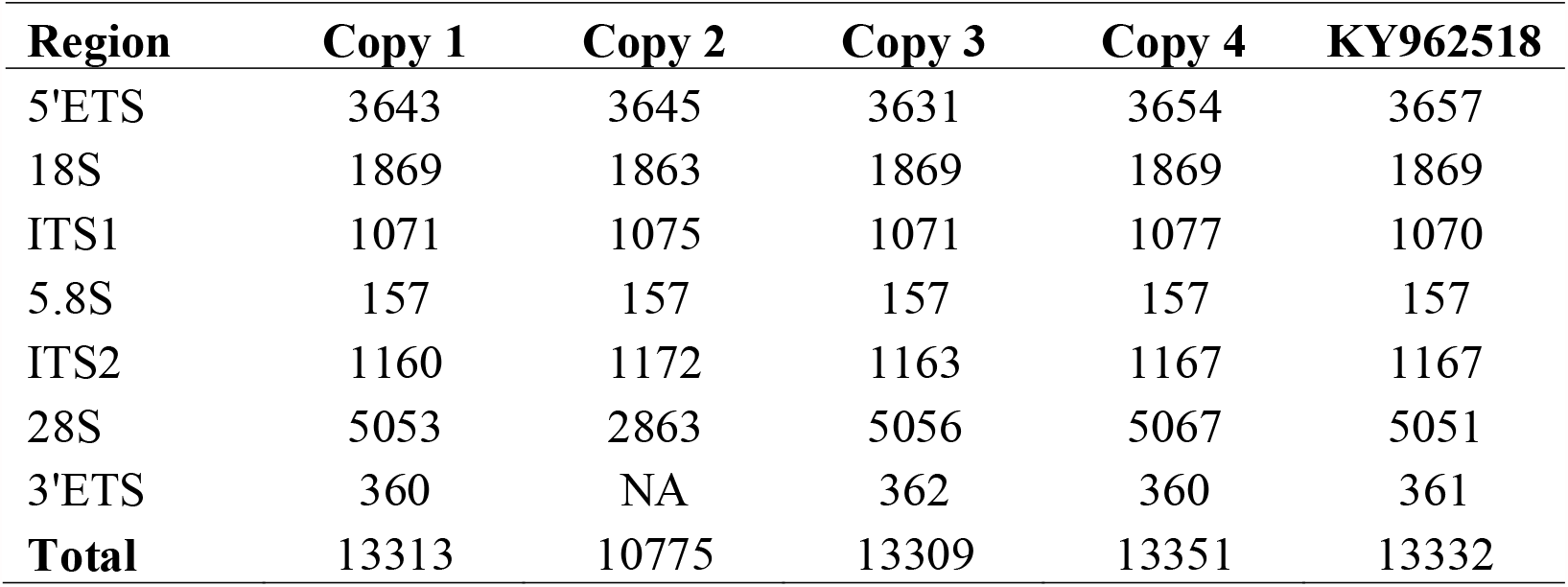
Coding region length (nt) variation of rDNA loci on chromosome 21.

**Table 2.** Variation in the rDNA loci on chromosome 21.

| **Region** | **SNV** | **INDEL** | **Total** |
| --- | --- | --- | --- |
| 5'ETS | 35 | 54 | 89 |
| 18S | 7 | 11 | 18 |
| ITS1 | 6 | 4 | 10 |
| 5.8S rRNA | 0 | 0 | 0 |
| ITS2 | 3 | 18 | 21 |
| 28S rRNA | 12 | 12 | 24 |
| 3'ETS | 0 | 1 | 1 |
| **Total** | 63 | 100 | 163 |

**Figure 1.**
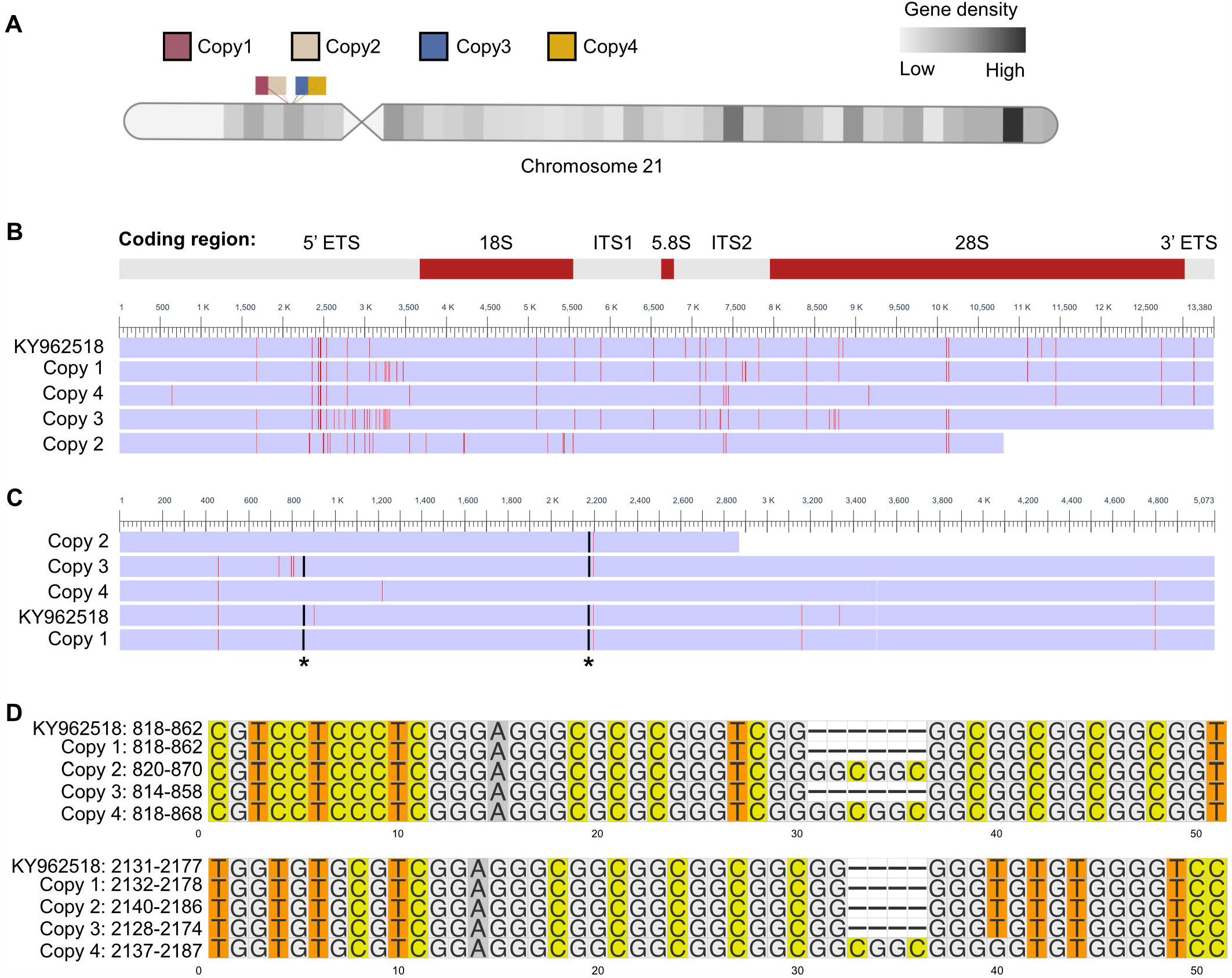
Multiple sequence alignment of human rRNA coding region in chromosome 21. (A) Schematic representation of four rDNA copies in human acrocentric chromosome 21 (GRCh38). rDNA copies are labeled by different colors. (B) Multisequence alignment of the coding regions of the chromosome 21 rDNA copies and KY962518 reference sequence. The differences are colored by red. rRNA coding regions are shown on the top. (C) Multisequence alignment of 28S rRNA. Asterisks indicate the position of small INDELs. (D) Multisequence alignment of two INDELs in 28S rRNA. Numbering on the left refers to 28S rRNA positions.

### High rDNA GC content anticorrelates with WGS read depth

The GC content of rDNA is high and reaches up to 90%. High genomic GC content poses a challenge to PCR-based sequencing technologies (Meienberg et al. 2015). To assess the impact of GC content on the sequencing depth throughout the rRNA coding region, we employed high coverage WGS data from the Cancer Cell Line Encyclopedia (CCLE) project (Barretina et al. 2012). For this purpose, we randomly selected 70 cancer cell lines, conducted read alignment to the reference KY962518 using Sentieon DNASeq tools and calculated the read depth across the rDNA sequence. We normalized the read depth per position against the total rDNA reads in the 13.3 kb coding region for each cancer cell line. The GC content was calculated in 70 base pair bins and correlated with the normalized read depth. This analysis showed that the GC content strongly negatively correlated with the read depth (Figure 2A). We then plotted the read depth and GC content per 70 bp bins. Notably, majority of the rDNA coding region read depth of the 70 cancer cell lines was high. Areas with very low read depths were evident, and largely correlated with areas of over 80% GC content (Figure 2B).

**Figure 2.**
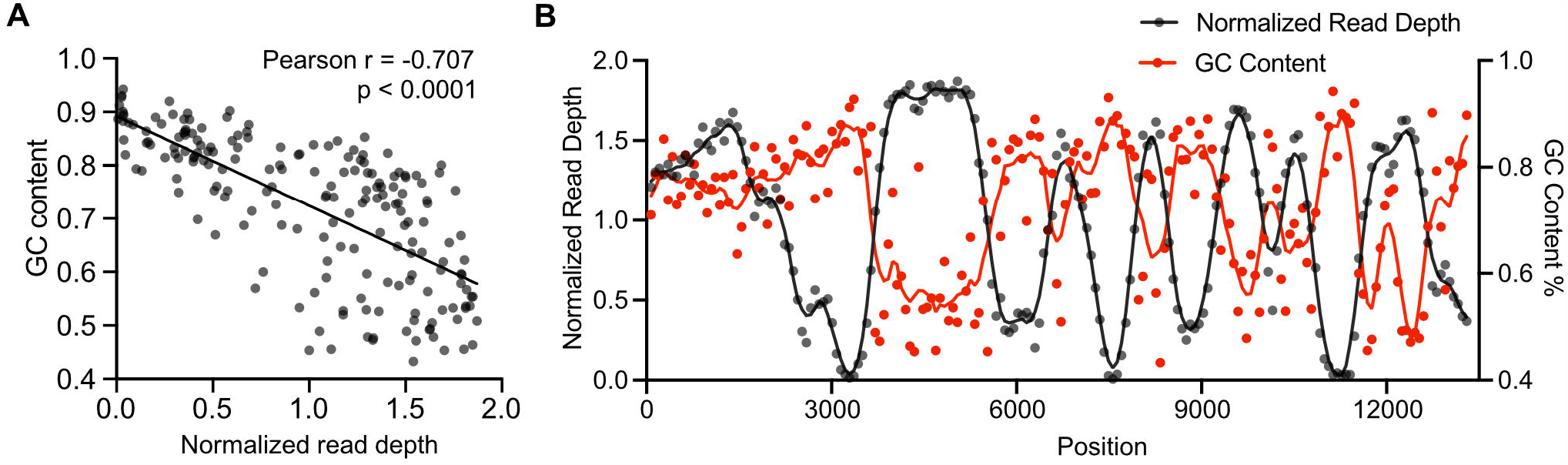
WGS sequence read depth anticorrelates with rDNA GC content. (A) Scatter plot showing GC content as compared to the normalized mean read depth in rDNA coding region. High coverage WGS data were obtained from 70 cancer cell lines in the CCLE database. *P* was determined by two-sided Pearson’s correlation test. (B) Normalized read depth and GC content are plotted across the rDNA coding region.

### Variant discovery in rRNA genes across human 2504 genomes

To explore rRNA gene variation in human genomes, we employed the 1000 Genomes Project high coverage WGS resource that has enabled variant discovery across populations (Genomes Project et al. 2015). We first examined the sequence coverage of rDNA and WGS data on 2,504 individuals using Samtools. Consistent with the description of the WGS data, we observed 30-fold or higher whole genome coverage of the 2,504 samples (Figure 3A). The coverage for rDNA was on average 5,000-fold in the human WGS dataset (Figure 3A). This estimation indicates that despite some highly challenging read areas exist, the coverage for the rDNA sequences is deep and suitable for variant calling.

**Figure 3.**
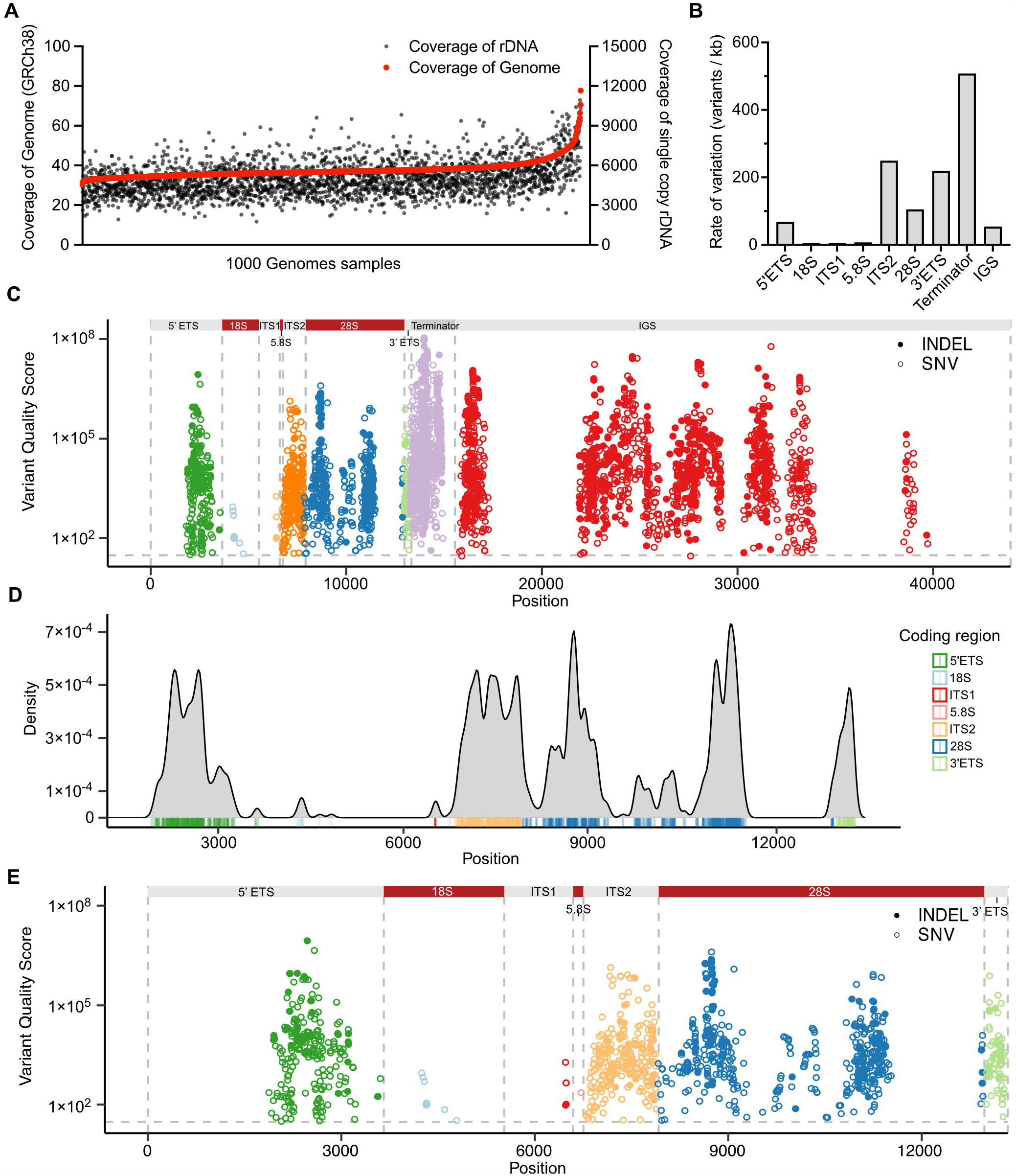
rRNA gene variant discovery in the human genome. (A) Coverage of rDNA and whole genome reads are shown for each individual from the 1000 Genomes Project (n = 2,504). (B) Number of variant positions in the rRNA gene domains. (C) Manhattan plot of variant distribution in a full-length 45 kb rDNA copy. Colors depict rRNA regions, which are indicated on the top. INDELs are show in solid dots, and SNVs in circles. Grey horizontal line represents the variant quality score significance (QUAL) threshold set at 30. (D) Density plot of the variants in the 13 kb rDNA coding region. Coding region domains are color-labeled as indicated by the legend. (E) Manhattan plot of variant distribution in the rRNA coding region. Color-coding as in C.

To perform large scale variant calling of WGS data, the standard software package for variant calling, GATK (Genome Analysis Toolkit) pipeline, is computing-intense for the analysis of single WGS samples even using multiple computing nodes and after optimization (Van der Auwera et al. 2013). To offer further computing speed over GATK, several ultrafast options have been developed (Raczy et al. 2013; Weber et al. 2016; Pluss et al. 2017). Given that the DNASeq pipeline of Sentieon provides faster variant calling without compromising accuracy, we employed it for the analysis of the genomic variation in human rDNA. The workflow was applied on WGS data of 2,504 individuals to perform variant calling against the four chromosome 21 rDNA copies, including the coding and IGS sequences, totaling 145 kb. Cut-off for variant calling was defined as variant quality score (QUAL) > 30 (< 0.1% error), and rare variants observed in < 3 alleles were excluded.

The final variant callset across the 2,504 samples identified 2,658 SNV and 1,132 INDEL positions (Table 3). The mean variant density across the rDNA coding region was 86 variants/kb. The density was highest on the terminator (over 500 variants/kb), whereas overall, was low on the IGS (54 variants/kb) (Figure 3B). The data showed that the terminator domain and IGS region had the largest number of variant positions (2,690 positions). Notably, the ITS2 and 3’ ETS domains had very high variant densities (over 200 variants/kb) (Figure 3B). The variants clustered around peaks with very high QUAL scores (> 10^6^) and positioned non-randomly on the rDNA (Figure 3C and Table 3).

**Table 3.**
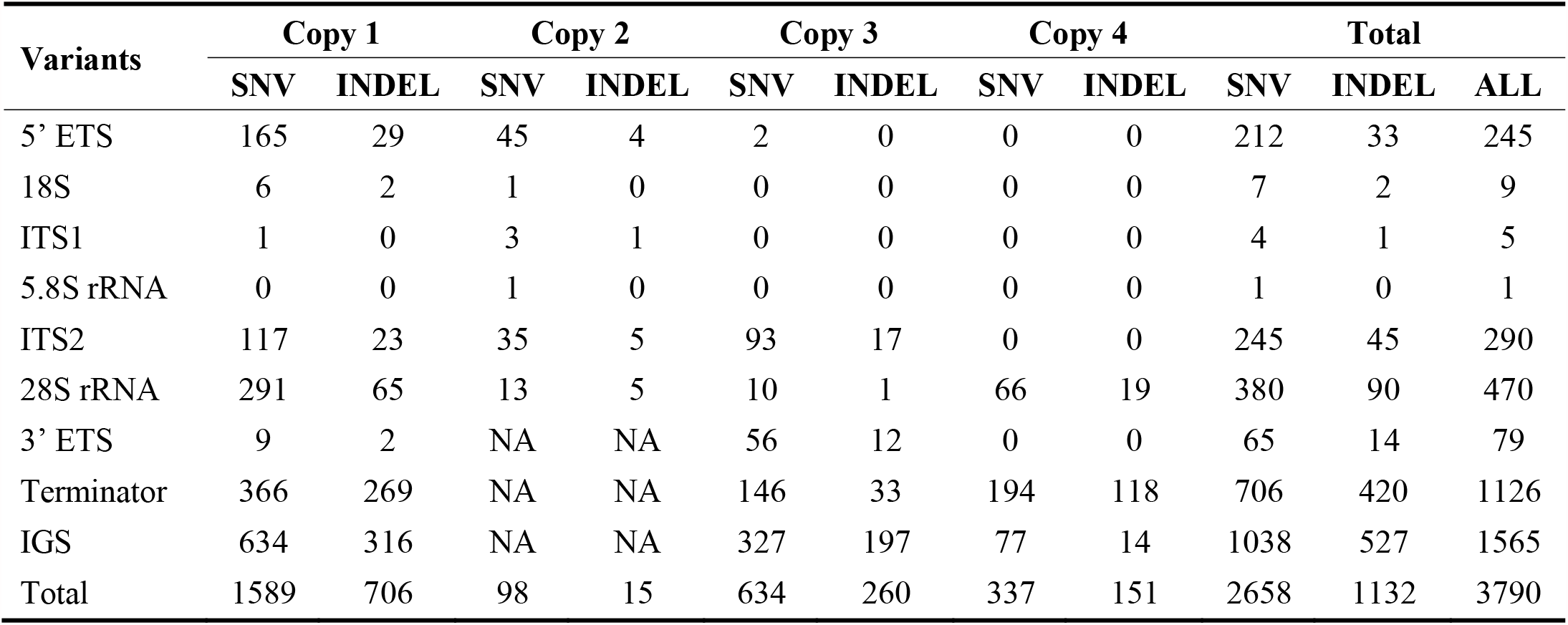
Variant calls of rDNA copies in 2,504 human genomes.

| **Variants** | **Copy 1** |  | **Copy 2** |  | **Copy 3** |  | **Copy 4** |  | **Total** |  |  |
| --- | --- | --- | --- | --- | --- | --- | --- | --- | --- | --- | --- |
|  | **SNV** | **INDEL** | **SNV** | **INDEL** | **SNV** | **INDEL** | **SNV** | **INDEL** | **SNV** | **INDEL** | **ALL** |
| 5' ETS | 165 | 29 | 45 | 4 | 2 | 0 | 0 | 0 | 212 | 33 | 245 |
| 18S | 6 | 2 | 1 | 0 | 0 | 0 | 0 | 0 | 7 | 2 | 9 |
| ITS1 | 1 | 0 | 3 | 1 | 0 | 0 | 0 | 0 | 4 | 1 | 5 |
| 5.8S rRNA | 0 | 0 | 1 | 0 | 0 | 0 | 0 | 0 | 1 | 0 | 1 |
| ITS2 | 117 | 23 | 35 | 5 | 93 | 17 | 0 | 0 | 245 | 45 | 290 |
| 28S rRNA | 291 | 65 | 13 | 5 | 10 | 1 | 66 | 19 | 380 | 90 | 470 |
| 3' ETS | 9 | 2 | NA | NA | 56 | 12 | 0 | 0 | 65 | 14 | 79 |
| Terminator | 366 | 269 | NA | NA | 146 | 33 | 194 | 118 | 706 | 420 | 1126 |
| IGS | 634 | 316 | NA | NA | 327 | 197 | 77 | 14 | 1038 | 527 | 1565 |
| Total | 1589 | 706 | 98 | 15 | 634 | 260 | 337 | 151 | 2658 | 1132 | 3790 |

We further visualized the variant density in the coding region (Figure 3D). In addition to the ITS2 and 3’ ETS domains, the 5’ ETS domain had a high density of variants clustered around 2,100 to 3,000 bp from transcription start site (Figure 3D). None of these variants position to the known human rRNA processing sites (Henras et al. 2015). Of note, 18S and 5.8S rRNA had only a few variants, whereas 470 variant positions and a total of 592 variants were observed in the 28S rRNA (Figure 3D and E). This indicates 18S and 5.8S rRNAs are highly conserved compared to 28S rRNA. These results reveal highly conserved and variable regions throughout the rDNA sequence and diversity across human genomes.

### Annotation of variants on the 28S rRNA structure

Given the abundance of variants in 28S rRNA and their clustering to distinct high and low density regions, and significance of the mature 28S rRNA for the 60S subunit function, we focused on annotation of the variants on 28S rRNA. We mapped the variant density profile on the human 28S rRNA secondary structure derived from Ribovision (Bernier et al. 2014). Interestingly, variants with the highest QUAL scores were located in the highly flexible expansion segment helical folds ES27L and ES7L (Figure 4A). To position the variants to the three-dimensional environment in the rRNA structure, we mapped the variant density profile to the 28S rRNA in the 80S ribosome cryo-EM structure (Anger et al. 2013). The regions with highest density of variants were located on the surface of the ribosome (Figure 4B). These regions represent expansion segments forming A-form helices that are under species-specific evolution (Anger et al. 2013; Fujii et al. 2018).

**Figure 4.**
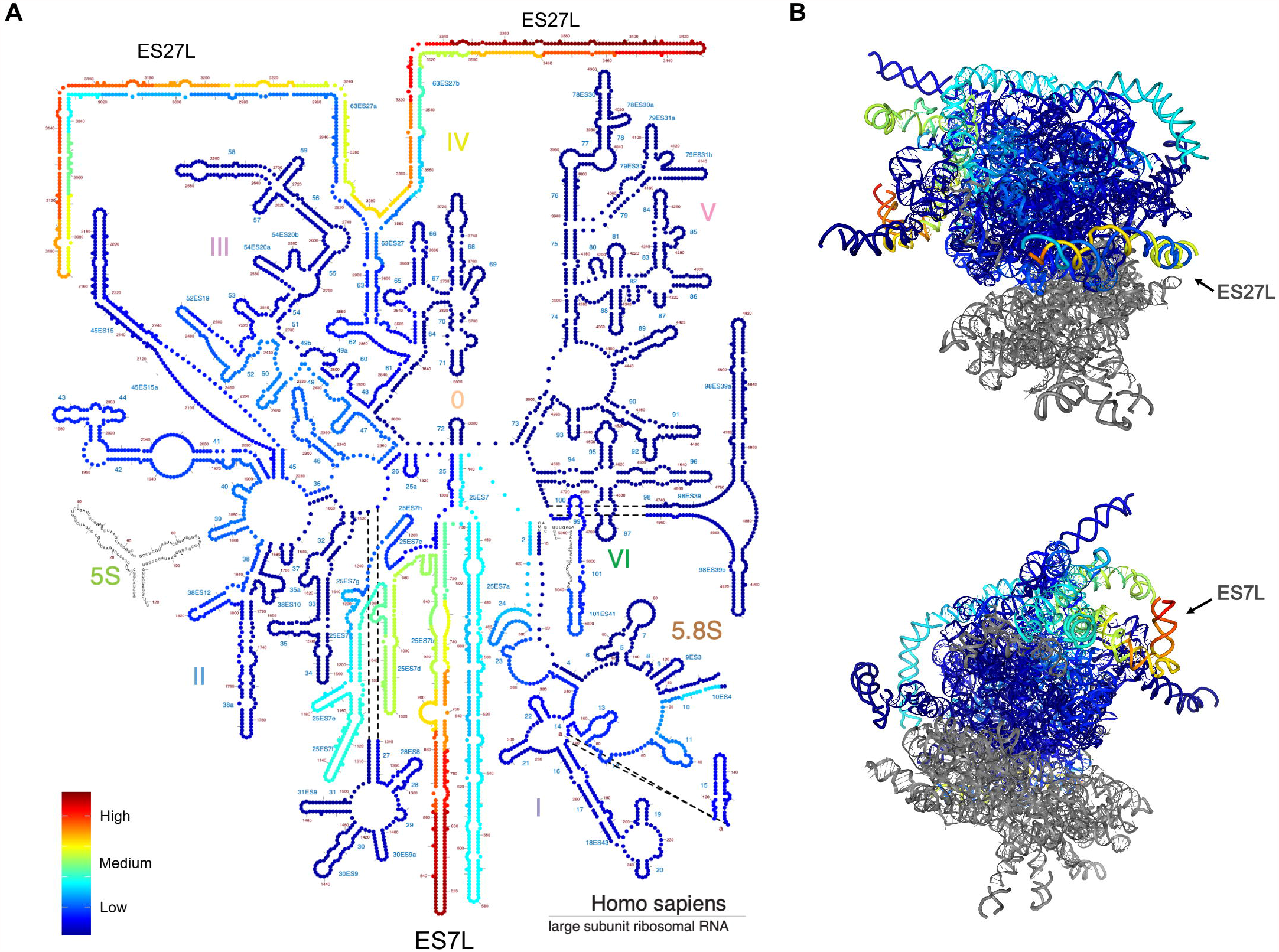
Annotation of variants on 28S rRNA structure. (A) Variant density on the secondary structure diagram of 28S rRNA. Nucleotides are color-labeled by the variant density value. (B) Variants of 28S rRNA (shown in *blue*) are shown on the 3D structure model (PDB 4V6X). Nucleotides are color-labeled by the variant density value as in A. High variant density expansion segments (ES) ES7L and ES27L are indicated. Images were generated with RiboVision.

Chemical modifications of human rRNA are introduced during ribosome biogenesis and required for the rRNA folding and stability. More than 130 individual rRNA modifications are indicated in the 3D structure of the human ribosome (Natchiar et al. 2017). We further compared the human genome variant positions to the recorded modification sites in 28S rRNA. Only one modification site (C2861 methylation) had a SNV in the human genomes, but had a very low allele count and QUAL score (7 and 99, respectively). The combined results suggest that the rRNA core ribosomal protein interaction sites and chemical modifications on rRNA are critical for functioning ribosomes and are rarely altered. The high number of variants in the 28S rRNA flexible expansion segments suggest that these may supply ribosome heterogeneity.

## DISCUSSION

The current knowledge of the variation in human rDNA sequences is incomplete. This study shows the heterogeneity of human rRNA genes across large populations. We show that diverse lengths and considerable number of variants exist within the established human rDNA GRCh38 chromosome 21 reference sequences. We annotated the human rDNA variants (SNV/INDEL) using the high coverage WGS data including 2,504 individuals from 26 ancestry-diverse populations. We performed SNV/INDEL calling on the GRCh38 reference rRNA gene copies and generated a comprehensive rDNA variant map. In total, 3,790 SNV/INDEL variant positions were identified. Density distribution analysis of variants showed noticeable clustering throughout the gene, including in the rRNA coding region, providing a perspective of both invariant and highly diverse regions. We mapped the 28S rRNA high density variants on the ribosome structure, which showed their location in the 28S rRNA expansion segments that are under continuous evolution. This suggests that the variation in 28S rRNA can potentially contribute to heterogeneity in ribosome function. The advantage of analysis of this large dataset, instead of single-genome sample comparisons, is that it not only affirms confidence of the reference genotype, but provides a resource to assess the impact of rRNA diversity in subsequent functional studies.

Genomic GC content affects sequencing depth due to PCR bias during the sequencing process (Dohm et al. 2008; Meienberg et al. 2015; Laursen et al. 2017). Reduction of GC bias is critical in improving the assembly of the genomes and increasing the accuracy of biomedical or clinical application. Given that the rDNA copies have high GC content regions in eukaryotes (Escobar et al. 2011; Parks et al. 2018) and that the PCR bias affects the accuracy of rDNA gene sequencing, we calculated the sequencing depth of rDNA using cancer cell line WGS data from the CCLE resource. Relatively low read depth was observed in the high GC content regions showing a significant negative correlation with the GC rich areas. Benefitting from the high coverage of 1000 genomes WGS data, we observe a 5000-fold coverage on rDNA region even in the high GC content regions enabling variant calling analyses. A high GC content will impact also other PCR-based methods such as chromatin and RNA immunoprecipitation and sequencing analyses, and has been observed in such studies as low read coverage areas. Computational methods and PCR-free technology in WGS have been developed to decrease this bias and to obtain better coverage of biologically important loci with high GC content (Benjamini and Speed 2012; Ross et al. 2013). Recently, the T2T consortium released the first complete human genome sequence that detailed the acrocentric rDNA arrays in the haploid CHM13 cells (Nurk et al. 2021). These advanced techniques and resources help in removing obstacles in identification of human variants both in discovery research and precision medicine.

Few studies have evaluated the heterogeneity of rRNA genes across human populations. This is partly due to the difficulty of assembling a reference for the highly variable rDNA loci. Unfortunately, assembly continues to be a computationally challenging problem for rDNA tandem repeat units using shotgun sequencing. A refined reference for a rDNA copy unit, assembled applying TAR cloning and long sequencing technologies has enabled further assessment of individual rDNA units on chromosomes 21 and 22 (Kim et al. 2018; Kim et al. 2021). Here, based on the refined human rDNA reference sequence (KY962518), we identified four copies of rDNA in the current human reference genome GRCh38 suitable for rDNA SNV/INDEL calling. Though they were 99% identical to the reference KY962518, each copy presented with both length and sequence variation, and a total of 163 variants were identified. This copy variation is consistent with the observation that chromosome 21 copies are mosaic (Nurk et al. 2021). Based on the T2T consortium data on CHM13 cell line HiFi reads, the degree of mosaicism varies between the acrocentric arrays being high on chromosomes 13, 15 and 21 and low on 14 and 22. The mosaicism in the copies used here is hence present as natural variation in the variant calling pipeline. The variant calling pipeline is designed to identify both intra-array and interindividual heterogeneity in the rRNA arrays compared to the reference. We took advantage of the 1000 Genomes Project, the largest fully open resource of whole genome sequencing data, making this resource an ideal starting point to identify human rDNA variants. Our analysis is based on the new high coverage WGS resource and generated a comprehensive set of rDNA SNV/INDELs.

In contrast to previous published studies, based on the earlier reference sequence U13369 (Babaian 2017; Xu et al. 2017; Parks et al. 2018), this study demonstrates distinct clustering of the variants to specific rRNA regions. The highest density of variants was observed in the rRNA transcription termination sites containing repetitive CT-sequences, as well as in the rRNA coding region 5’ ETS and ITS2 domains. However, even in these domains no variants were detected in their known rRNA processing sites. Also, the 18S and 5.8S rRNA coding sequences, in agreement with Xu et al. 2017, but in contrast to Parks et al., 2018, were near invariant in this deep dataset, suggesting a low tolerance for further adaptation of these mature rRNA coding sequences. Also, in contrast to the study by Parks et al., 2018 we did not detect variants in the 28S rRNA modification sites or high-confidence, frequent variants in the ribosome bridge intersubunit sites. Notably, and surprisingly, large areas of the IGS were devoid of variants. This is in contrast to the earlier notion that IGS is highly variable (Gonzalez and Sylvester 1995; Gonzalez and Sylvester 2001). The conservation of IGS may be driven by as of yet unidentified functional relevance of these sequences. Stress-inducible non-coding IGS-derived RNAs have been identified, and IGS regions with transcriptionally active chromatin states, as well as both sense and antisense transcription by Pol I and Pol II (Audas et al. 2012; Agrawal and Ganley 2018; Abraham et al. 2020). On the other hand, high variant densities with high QUAL scores were observed in the IGS at 16, 23-24, 28, 31, 33 and 39 kb that frequently were represented by CT repeat and TG repeat sequences. Overall, these results suggest a high degree of conservation of not only the rDNA arrays between chromosomes, but also low variance between individuals. Given that the rDNA copies are considered to undergo frequent genomic alterations, this conservation is remarkable.

Genomic variants of rRNA genes may provide a molecular basis for physically and functionally heterogeneous ribosomes. rDNA unit diversity has been observed in various species (Tseng et al. 2008; Matyasek et al. 2012; Agrawal and Ganley 2018). In mouse, rDNA array consists of genetically distinct variants, and a subset of rDNA genes are regulated in a cell-type-specific manner (Tseng et al. 2008). Mouse rRNA variants are differentially expressed in organs and the epigenetic state of rDNA copies is influenced by *in utero* nutrition (Holland et al. 2016; Parks et al. 2018). Distinct composition and functionality of ribosomes has been proposed within eukaryotic cells (Sauert et al. 2015; Shi and Barna 2015; Slavov et al. 2015; Briggs and Dinman 2017; Mills and Green 2017; Emmott et al. 2019). Ribosome-associated proteins further functionally diversify mammalian ribosomes (Simsek et al. 2017). Heterogeneous ribosomes, containing diverse post translational modifications of the ribosomal proteins or distinct ribosome binding factors, can preferentially translate different subsets of mRNAs (Emmott et al. 2019; Li and Wang 2020). However, whether variation in the rRNA coding sequences have an impact on ribosomal functional heterogeneity remains poorly understood. The expansion segments, especially ES27L, are dynamic and assist in connecting the 40S mRNA exit and 60S tunnel exit sites on the ribosome, and interact with export factors and other regulators (Anger et al. 2013). For example, the human ES27L acts as an RNA scaffold to facilitate binding of the methionine amino peptidase (MetAP) to control translation fidelity (Fujii et al. 2018). We observed that positions relevant for the rRNA-ribosomal protein interaction are much less variant compared to the highly flexible human-expanded rRNA ES7L and ES27L helical folds on the surface of complex. Hence, variation in 28S ES27L may provide a genetic basis for heterogeneous translation fidelity in human individuals. On the other hand, our data show that rRNA modification sites are highly conserved. This is consistent with the notion that rRNA-ribosomal protein interaction sites and chemical modifications on rRNA are critical for ribosome assembly and function and under strict preservation. Yet, the high number of 28S rRNA variants implicates that they potentially contribute to translation control or mediate selective mRNA translation. However, this study does not account for putative tissue-specific heterogeneity or somatic alterations of the rDNA genes.

In addition to the challenge of the sequence identity of the gene copies, the rDNA repeats are highly dynamic due to recombination events during meiosis, DNA repair and in diseases, such as cancer. Identification of disease-linked variants will require comprehensive assessment of the rDNA variation in the normal human genomes. Our study of genetic variation on the rRNA genes across human populations, aided by high coverage WGS data, narrows down the previously presumed variation and provides a resource of rRNA variant detection and basis for the understanding of the impact of rRNA variants potentially affecting physiology and disease.

## MATERIALS AND METHODS

### Identification of rDNA regions in the human reference genome GRCh38

rDNA reference KY962518 (GenBank: KY962518.1) was aligned to the human genome reference genome (*Homo sapiens* genome assembly GRCh38.p13) by using Basic Local Alignment Search Tool (BLAST). Highly similar sequences (Megablast) with 100% query coverage and ≥ 99% percent identity were considered as high confidence rDNA copies. Unlocalized genomic scaffold and PATCHES sequence were excluded from blast results. rDNA locus on chromosome 21 was visualized using RIdeogram R package (Hao et al. 2020).

### Multiple sequence alignments

Multiple sequence alignment of the rDNA reference KY962518 and rDNA copies identified on chromosome 21 was performed with Clustal Omega Multiple Sequence Alignment (MSA) to detect variation across copies. The alignment results were visualized by NCBI MSA viewer. Number of SNVs and INDELs were counted by Jvarkit and VCFtools (Danecek et al. 2011). INDELs from multiple sequence alignment were visualized using ggmsa R package (Yu 2020).

### Comparison of GC content and read depth

Genomic GC content percentage was calculated as Count(G + C) / Count(A + T + G + C) * 100%. A GC% value was computed for each 70 nucleotide interval in rDNA reference KY962518. We randomly selected 70 cancer cell lines from CCLE project, downloaded related WGS data and performed read alignment to KY962518 using Sentieon DNASeq tools. Read depth at each position or region was computed by samtools depth according to Samtools manual. For each sample, we normalized the read depth per position as depth per position/mean depth in the rDNA coding region. Mean normalized read depth values for 70 cancer cell lines are presented. We then calculated Pearson correlation between the GC content and mean normalized read depth of the 70 samples.

### Coverage for rDNA and WGS sequence reads

The average rDNA read coverage was computed by samtools flagstat according to protocol in Samtools manual for WGS read analysis. GRCh38 was used as reference and reads mapping to the identified rDNA copies were calculated. Mapped reads for each sample were extracted and rDNA coverage was computed as (mapped read count * read length) / rDNA copy size. Genome sequence read coverage was computed as (mapped read count * read length) / total genome size.

### Variant calling and variant density measurements

Hight coverage WGS data for 2,504 individuals was obtained from the 1000 genome project and read alignment to the human reference genome GRCh38, duplicate marking, and Base Quality Score Recalibration (BQSR) was performed (Byrska-Bishop et al. 2021). Alignment files in CRAM format were used for variant calling. Germline SNVs and INDEL variants were called with the DNAseq pipeline (Sentieon, Release 201911). We used the following variant call criteria: QUAL score >30 PHRED (p < 0.001) and presence in three or more alleles. Computing was conducted using Maryland Advanced Research Computing Center (MARCC) high performance cluster computing nodes. Integrative Genomics Viewer (IGV, version 2.8.6) was used to visualize read alignments. SNVs and INDELs were summarized from gVCF files derived by Sentieon DNAseq pipeline and visualized by ggplot2 R package. The density of variants located in rDNA region was calculated and visualized by ggridges and ggplot2 R package . Bandwidth used for density calculation was provided as 50.

### Annotation of variants on 28S rRNA 2D and 3D structure

Ribosome Visualization Suite RiboVision2 (http://apollo.chemistry.gatech.edu/RiboVision2/) was used to visualize the density of variants on 28S rRNA (Bernier et al. 2014). Variant density was mapped on the secondary (2D) structure of 28S rRNA and portrayed by color gradient. All proteins in the tool were checked for visualizing the protein contacts on 28S rRNA. The variant densities were mapped to the three-dimensional (3D) structures of ribosomes and the density intensity is indicated by a color gradient. Images were generated using RiboVision2 based on PDB 4V6X (Anger et al. 2013).

## ACKNOWLEDGEMENTS

This work was supported in part by NIGMS R01GM121404 and NCI P30 CA006973. We thank Dr. Steven Salzberg for helpful discussions and Maryland Advanced Research Computing Center (MARCC) for access to the high computing resource.

## SUPPLEMENTAL MATERIAL

**Supplemental Table 1.** rDNA loci positions and similarities on chromosome 21.

| **Copy** | **Copy start** | **Copy stop** | **Length** | **% Identity** |
| --- | --- | --- | --- | --- |
| 1 | 8205989 | 8219303 | 13313 | 99.8 |
| 2 | 8250198 | 8260973 | 10775 | 99.0 |
| 3 | 8389036 | 8402344 | 13309 | 99.4 |
| 4 | 8433222 | 8446572 | 13351 | 99.6 |

**Supplemental Table 2**. rDNA variant calls in the human genomes. [separate file]

## REFERENCES

Abraham KJ, Khosraviani N, Chan JNY, Gorthi A, Samman A, Zhao DY, Wang M, Bokros M, Vidya E, Ostrowski LA et al. 2020. Nucleolar RNA polymerase II drives ribosome biogenesis. Nature 585: 298–302.

Agrawal S, Ganley ARD. 2018. The conservation landscape of the human ribosomal RNA gene repeats. PLoS One 13: e0207531.

Anger AM, Armache JP, Berninghausen O, Habeck M, Subklewe M, Wilson DN, Beckmann R. 2013. Structures of the human and Drosophila 80S ribosome. Nature 497: 80–85.

Audas TE, Jacob MD, Lee S. 2012. Immobilization of proteins in the nucleolus by ribosomal intergenic spacer noncoding RNA. Mol Cell 45: 147–157.

Babaian A. 2017. Intra-and Inter-individual genetic variation in human ribosomal RNAs. BioRxiv: 118760.

Barretina J, Caponigro G, Stransky N, Venkatesan K, Margolin AA, Kim S, Wilson CJ, Lehár J, Kryukov GV, Sonkin D. 2012. The Cancer Cell Line Encyclopedia enables predictive modelling of anticancer drug sensitivity. Nature 483: 603–607.

Benjamini Y, Speed TP. 2012. Summarizing and correcting the GC content bias in high-throughput sequencing. Nucleic Acids Res 40: e72.

Bernier CR, Petrov AS, Waterbury CC, Jett J, Li F, Freil LE, Xiong X, Wang L, Migliozzi BL, Hershkovits E. 2014. RiboVision suite for visualization and analysis of ribosomes. Faraday discussions 169: 195–207.

Briggs JW, Dinman JD. 2017. Subtractional Heterogeneity: A Crucial Step toward Defining Specialized Ribosomes. Mol Cell 67: 3–4.

Byrska-Bishop M, Evani US, Zhao X, Basile AO, Abel HJ, Regier AA, Corvelo A, Clarke WE, Musunuri R, Nagulapalli K et al. 2021. High coverage whole genome sequencing of the expanded 1000 Genomes Project cohort including 602 trios. bioRxiv: 2021.2002.2006.430068.

Danecek P, Auton A, Abecasis G, Albers CA, Banks E, DePristo MA, Handsaker RE, Lunter G, Marth GT, Sherry ST et al. 2011. The variant call format and VCFtools. Bioinformatics 27: 2156–2158.

Dohm JC, Lottaz C, Borodina T, Himmelbauer H. 2008. Substantial biases in ultra-short read data sets from high-throughput DNA sequencing. Nucleic Acids Res 36: e105.

Emmott E, Jovanovic M, Slavov N. 2019. Ribosome Stoichiometry: From Form to Function. Trends Biochem Sci 44: 95–109.

Escobar JS, Glemin S, Galtier N. 2011. GC-biased gene conversion impacts ribosomal DNA evolution in vertebrates, angiosperms, and other eukaryotes. Mol Biol Evol 28: 2561–2575.

Fujii K, Susanto TT, Saurabh S, Barna M. 2018. Decoding the Function of Expansion Segments in Ribosomes. Mol Cell 72: 1013–1020 e1016.

Genomes Project C, Auton A, Brooks LD, Durbin RM, Garrison EP, Kang HM, Korbel JO, Marchini JL, McCarthy S, McVean GA et al. 2015. A global reference for human genetic variation. Nature 526: 68–74.

Gonzalez IL, Gorski JL, Campen TJ, Dorney D, Erickson JM, Sylvester JE, Schmickel RD. 1985. Variation among human 28S ribosomal RNA genes. Proceedings of the National Academy of Sciences 82: 7666–7670.

Gonzalez IL, Sylvester JE. 1995. Complete sequence of the 43-kb human ribosomal DNA repeat: analysis of the intergenic spacer. Genomics 27: 320–328.

Gonzalez IL, Sylvester JE. 2001. Human rDNA: evolutionary patterns within the genes and tandem arrays derived from multiple chromosomes. Genomics 73: 255–263.

Gonzalez IL, Sylvester JE, Schmickel RD. 1988. Human 28S ribosomal RNA sequence heterogeneity. Nucleic Acids Res 16: 10213–10224.

Hao Z, Lv D, Ge Y, Shi J, Weijers D, Yu G, Chen J. 2020. RIdeogram: drawing SVG graphics to visualize and map genome-wide data on the idiograms. PeerJ Comput Sci 6: e251.

Henderson A, Warburton D, Atwood K. 1972. Location of ribosomal DNA in the human chromosome complement. Proceedings of the National Academy of Sciences 69: 3394–3398.

Henras AK, Plisson-Chastang C, O’Donohue MF, Chakraborty A, Gleizes PE. 2015. An overview of pre-ribosomal RNA processing in eukaryotes. Wiley Interdiscip Rev RNA 6: 225–242.

Holland ML, Lowe R, Caton PW, Gemma C, Carbajosa G, Danson AF, Carpenter AA, Loche E, Ozanne SE, Rakyan VK. 2016. Early-life nutrition modulates the epigenetic state of specific rDNA genetic variants in mice. Science 353: 495–498.

Khatter H, Myasnikov AG, Natchiar SK, Klaholz BP. 2015. Structure of the human 80S ribosome. Nature 520: 640–645.

Kim JH, Dilthey AT, Nagaraja R, Lee HS, Koren S, Dudekula D, Wood Iii WH, Piao Y, Ogurtsov AY, Utani K et al. 2018. Variation in human chromosome 21 ribosomal RNA genes characterized by TAR cloning and long-read sequencing. Nucleic Acids Res 46: 6712–6725.

Kim JH, Noskov VN, Ogurtsov AY, Nagaraja R, Petrov N, Liskovykh M, Walenz BP, Lee HS, Kouprina N, Phillippy AM et al. 2021. The genomic structure of a human chromosome 22 nucleolar organizer region determined by TAR cloning. Sci Rep 11: 2997.

Kuo BA, Gonzalez IL, Gillespie DA, Sylvester JE. 1996. Human ribosomal RNA variants from a single individual and their expression in different tissues. Nucleic acids research 24: 4817–4824.

Laursen MF, Dalgaard MD, Bahl MI. 2017. Genomic GC-Content Affects the Accuracy of 16S rRNA Gene Sequencing Based Microbial Profiling due to PCR Bias. Front Microbiol 8: 1934.

Li D, Wang J. 2020. Ribosome heterogeneity in stem cells and development. J Cell Biol 219.

Matyasek R, Renny-Byfield S, Fulnecek J, Macas J, Grandbastien MA, Nichols R, Leitch A, Kovarik A. 2012. Next generation sequencing analysis reveals a relationship between rDNA unit diversity and locus number in Nicotiana diploids. BMC Genomics 13: 722.

McStay B. 2016. Nucleolar organizer regions: genomic ‘dark matter’ requiring illumination. Genes Dev 30: 1598–1610.

Meienberg J, Zerjavic K, Keller I, Okoniewski M, Patrignani A, Ludin K, Xu Z, Steinmann B, Carrel T, Rothlisberger B et al. 2015. New insights into the performance of human whole-exome capture platforms. Nucleic Acids Res 43: e76.

Mills EW, Green R. 2017. Ribosomopathies: There’s strength in numbers. Science 358: eaan2755.

Moss T, Langlois F, Gagnon-Kugler T, Stefanovsky V. 2007. A housekeeper with power of attorney: the rRNA genes in ribosome biogenesis. Cell Mol Life Sci 64: 29–49.

Natchiar SK, Myasnikov AG, Kratzat H, Hazemann I, Klaholz BP. 2017. Visualization of chemical modifications in the human 80S ribosome structure. Nature 551: 472–477.

Nurk S, Koren S, Rhie A, Rautiainen M, Bzikadze AV, Mikheenko A, Vollger MR, Altemose N, Uralsky L, Gershman A et al. 2021. The complete sequence of a human genome. bioRxiv: 2021.2005.2026.445798.

Parks MM, Kurylo CM, Dass RA, Bojmar L, Lyden D, Vincent CT, Blanchard SC. 2018. Variant ribosomal RNA alleles are conserved and exhibit tissue-specific expression. Sci Adv 4: eaao0665.

Pluss M, Kopps AM, Keller I, Meienberg J, Caspar SM, Dubacher N, Bruggmann R, Vogel M, Matyas G. 2017. Need for speed in accurate whole-genome data analysis: GENALICE MAP challenges BWA/GATK more than PEMapper/PECaller and Isaac. Proc Natl Acad Sci U S A 114: E8320–E8322.

Raczy C, Petrovski R, Saunders CT, Chorny I, Kruglyak S, Margulies EH, Chuang HY, Kallberg M, Kumar SA, Liao A et al. 2013. Isaac: ultra-fast whole-genome secondary analysis on Illumina sequencing platforms. Bioinformatics 29: 2041–2043.

Ross MG, Russ C, Costello M, Hollinger A, Lennon NJ, Hegarty R, Nusbaum C, Jaffe DB. 2013. Characterizing and measuring bias in sequence data. Genome Biol 14: R51.

Sauert M, Temmel H, Moll I. 2015. Heterogeneity of the translational machinery: Variations on a common theme. Biochimie 114: 39–47.

Shi Z, Barna M. 2015. Translating the genome in time and space: specialized ribosomes, RNA regulons, and RNA-binding proteins. Annu Rev Cell Dev Biol 31: 31–54.

Simsek D, Tiu GC, Flynn RA, Byeon GW, Leppek K, Xu AF, Chang HY, Barna M. 2017. The Mammalian Ribo-interactome Reveals Ribosome Functional Diversity and Heterogeneity. Cell 169: 1051–1065 e1018.

Slavov N, Semrau S, Airoldi E, Budnik B, van Oudenaarden A. 2015. Differential stoichiometry among core ribosomal proteins. Cell reports 13: 865–873.

Sloan KE, Warda AS, Sharma S, Entian KD, Lafontaine DLJ, Bohnsack MT. 2017. Tuning the ribosome: The influence of rRNA modification on eukaryotic ribosome biogenesis and function. RNA Biol 14: 1138–1152.

Tseng H, Chou W, Wang J, Zhang X, Zhang S, Schultz RM. 2008. Mouse ribosomal RNA genes contain multiple differentially regulated variants. PLoS One 3: e1843.

Van der Auwera GA, Carneiro MO, Hartl C, Poplin R, Del Angel G, Levy-Moonshine A, Jordan T, Shakir K, Roazen D, Thibault J. 2013. From FastQ data to high-confidence variant calls: the genome analysis toolkit best practices pipeline. Current protocols in bioinformatics 43: 11.10. 11-11.10. 33.

van Sluis M, van Vuuren C, Mangan H, McStay B. 2020. NORs on human acrocentric chromosome p-arms are active by default and can associate with nucleoli independently of rDNA. Proc Natl Acad Sci U S A 117: 10368–10377.

Weber JA, Aldana R, Gallagher BD, Edwards JS. 2016. Sentieon DNA pipeline for variant detection-Software-only solution, over 20× faster than GATK 3.3 with identical results. PeerJ PrePrints 4: e1672v1672.

Worton RG, Sutherland J, Sylvester JE, Willard HF, Bodrug S, Dube I, Duff C, Kean V, Ray PN, Schmickel RD. 1988. Human ribosomal RNA genes: orientation of the tandem array and conservation of the 5’ end. Science 239: 64–68.

Xu B, Li H, Perry JM, Singh VP, Unruh J, Yu Z, Zakari M, McDowell W, Li L, Gerton JL. 2017. Ribosomal DNA copy number loss and sequence variation in cancer. PLoS Genet 13: e1006771.

Yu G. 2020. Using ggtree to visualize data on tree-like structures. Current protocols in bioinformatics 69: e96.

